# Functional diversity and evolution of the *Drosophila* sperm proteome

**DOI:** 10.1101/2022.02.14.480191

**Authors:** Martin D. Garlovsky, Jessica Sandler, Timothy L. Karr

**Affiliations:** Department of Applied Zoology, Faculty of Biology, Technische Universität Dresden, Dresden 01069, Germany; Biosciences Mass Spectrometry Core Research Facility, Knowledge Enterprise, Arizona State University, USA; Neurodegenerative Disease Research Center, The Biodesign Institute, Arizona State University, USA

**Keywords:** Spermatozoa, seminal fluid proteins, ribosomes, meiotic sex chromosome inactivation, fertility, evolution, discovery proteomics, human disease, OMIM, *Drosophila*

## Abstract

Given the central role fertilization plays in the health and fitness of sexually reproducing organisms and the well-known evolutionary consequences of sexual selection and sperm competition, knowledge gained by a deeper understanding of sperm (and associated reproductive tissues) proteomes has proven critical to the field’s advancement. Due to their extraordinary complexity, proteome depth-of-coverage is dependent on advancements in technology and related bioinformatics, both of which have made significant advancements in the decade since the last *Drosophila* sperm proteome was published. Here we provide an updated version of the *Drosophila melanogaster* sperm proteome (DmSP3) using improved separation and detection methods and an updated genome annotation. We identified 2563 proteins, with label-free quantitation (LFQ) for 2125 proteins. Combined with previous versions of the sperm proteome, the DmSP3 contains a total of 3176 proteins. The top 20 most abundant proteins contained the structural elements *α*- and *β*-tubulins and sperm leucyl-aminopeptidases (S-Laps). Both gene content and protein abundance were significantly reduced on the X chromosome, a finding consistent with prior genomic studies of the X chromosome gene content and evolution. We identified 9 of the 16 Y-linked proteins, including known testis-specific male fertility factors. LFQ measured significant levels for 75/83 ribosomal proteins (RPs) we identified, including a number of core constituents. The role of this unique subset of RPs in sperm is unknown. Surprisingly, our expanded sperm proteome also identified 122 seminal fluid proteins (Sfps), proteins found predominantly in the accessory glands. The possibility of tissue contamination from seminal vesicle or other reproductive tissues was addressed using concentrated salt and detergent treatments. Salt treatment had little effect on sperm proteome composition suggesting only minor contamination during sperm isolation while a significant fraction of Sfps remained associated with sperm following detergent treatment suggesting Sfps may arise within, and have additional functions, in sperm *per se*.

## Introduction

Spermatozoa form, function and evolution is determined in large measure by its proteome (1). High throughput proteomics using liquid-chromatography tandem massspectrometry (LC-MS) has now been used to characterize the composition of the sperm proteome in a wide range of animals (1–3). These studies have revealed several common features of sperm as expected for an ancient cell type with a highly conserved function despite exhibiting exceptional morphological diversity across the tree of life (4–6). For instance, across taxa, sperm show enrichment of metabolic processes, mitochondria, axoneme, microtubules and cytoskeletal components (2, 3, 7, 8).

Recent advances in LC-MS technology, particularly in data acquisition time and improved liquid chromatographic systems, provide enhanced proteome coverage of complex cell and tissue types (9, 10). These advances accordingly allow routine and accurate quantitation of both label- and label-free methodologies, an essential element for comparative studies of sperm composition and function (11, 12). Additionally, these advances permit direct injection of sample peptides without the need for pre-fractionation using polyacrylamide gel electrophoresis thus avoiding sample loss. Accordingly, in the current study we re-interrogated the *Drosophila melanogaster* sperm proteome using direct solubilization of sperm followed by on-line fractionation of tryptic peptides and report on a significant increase in both proteome size and content.

*D. melanogaster* with its excellent genome annotation provides a powerful genetic and functional genomics model system to understand reproduction and fertility (e.g., (13)). Our previous efforts identified over 1,000 *D. melanogaster* sperm proteins with prior versions designated DmSP1 (7) and DmSP2 (14). The DmSP3 described in this study significantly increases coverage and refinement of the *D. melanogaster* sperm proteome, from the 1108 sperm proteins identified in the combined DmSP1 and DmSP2 (14) to more than 3000 proteins in the DmSP3 (Table 1). Table 1 highlights our extended knowledge base not only in terms of absolute numbers of sperm proteins, but also discovery of new protein groups including the surprising findings of substantial numbers of seminal fluid proteins and ribosomal proteins in the DmSP3. We use the increased proteome coverage and quantitative LFQ information in the DmSP3 to provide a detailed analysis of relative abundance of sperm proteins for the first time, and re-examine the evolutionary dynamics, gene age, and chromosomal distribution of proteins in the DmSP3. The analyses provide stronger support for previous claims and in particular cements the subjective prior findings supporting the meiotic sex-chromosome inactivation model for male-specific gene and X chromosome evolution (15–18).

**Table 1.** History of the *Drosophila melanogaster* sperm proteome (DmSP). DmSP1: (7); DmSP2: (14); DmSP3: this study. The DmSP2 combined the 341 proteins identified in the DmSP1 with the 956 proteins identified in the DmSP2. Likewise, the DmSP3 reported here represents the combined total of all proteins identified in the DmSP2 (n = 1108) with the 2563 proteins identified across all experiments in the current study. Numbers in parentheses denote number of newly identified proteins. *Under = significant gene underrepresentation compared to expected value (see Methods); ns = not significant.

|  | DmSP1 | DmSP2 | DmSP3 |
| --- | --- | --- | --- |
| Methods/Technology | LC-MS<sup>2</sup>/Maldi | SDS-PAGE | Cell digest |
| Machine | Thermo LCQ | LTQ Orbitrap | Orbitrap Fusion Lumos |
| Proteins identified | 341 | 1108 (+767) | 3176 (+2068) |
| X-linked* | Under | ns | Under |
| Y-linked | - | 4 | 9 (+5) |
| Sfps | CG2918 | 11 (+10) | 122 (+111) |
| Ribosomal proteins | - | 9 | 83 (+74) |

## Methods

### A. Fly stocks and sample preparation

We used laboratory wild-type strain Oregon-R *D. melanogaster* males, aged 5-7 days. All dissections tissue removal and sperm isolation were performed at room temperature in freshly prepared phosphate buffered saline (PBS) with or without protease inhibitors (HALT, Thermo Fisher). We anaesthetized flies and removed reproductive tracts with forceps under a stereo dissecting microscope as previously described (14). Briefly, each biological replicate from 10 males (20 paired seminal vesicles) were prepared separately over the course of no more than one hour by first removing the seminal vesicles from each male reproductive tract (containing testes, seminal vesicles, and accessory glands) into a fresh drop of PBS. Sperm were then carefully removed using fine needles to a 1.5ml microcentrifuge tube containing PBS (on ice). Sperm were then pelleted at 15,000 rpm for 15 minutes at 4°C, washed 3X with PBS and immediately solubilized in 25 microliters of 5% SDS/50mM TEAB containing 50mM dithiothreitol. Solubilized samples were then incubated for 10-15 minutes at 95°C and spun again at 15,000 rpm for 15 minutes at 20°C. No visible pellets were observed and the supernatants removed and stored at −20°C or immediately processed as described below.

Solubilized sperm proteins were quantified using EZQ Protein Quantitation Kit (Thermo Fisher) and 2.25ug alkylated (Pierce) using 40mM final concentration freshly prepared iodoacetamide for 30 minutes in the dark at room temperature. Samples were processed using the Protifi S-trap Micro Columns and instructions were given via the S-trap Ultra High Recovery Protocol (Protifi). Briefly, samples were acidified by addition of 12% phosphoric acid to a final concentration of~1.2% phosphoric acid. Proteins were digested by addition of 2.0 μg of porcine trypsin (MS grade, Pierce) and incubated at 30°C for 2 hours. S-trap buffer (90% methanol, 100 mM TEAB final) was also added in volumes 7X our total sample volume. Acidified sample and the S-trap buffer was filtered through columns. Columns were washed 3X with Strap buffer. An additional 0.5 μg of trypsin and 25 μL of 50 mM TEAB was added to the top of each column and incubated for 1 hour at 47°C. Samples were eluted off the S-trap columns using three elution buffers: 50 mM TEAB, 0.2% formic acid in water, and 50% acetonitrile/50% water + 0.2% formic acid. Samples were dried down via speed vac and resuspended in 20-30 μL of 0.1% formic acid.

### B. Liquid-chromatography tandem mass-spectrometry

All LC-MS analyses were performed at the Biosciences Mass Spectrometry Core Facility (https://cores.research.asu.edu/mass-spec/) at Arizona State University. All data-dependent mass spectra were collected in positive mode using an Orbitrap Fusion Lumos mass spectrometer (Thermo Scientific) coupled with an UltiMate 3000 UHPLC (Thermo Scientific). One μL of peptides were fractionated using an Easy-Spray LC column (50 cm × 75 μm ID, PepMap C18, 2 μm particles, 100 Å pore size, Thermo Scientific) equipped with an upstream 300um x 5mm trap column. Electrospray potential was set to 1.6 kV and the ion transfer tube temperature to 300°C. The mass spectra were collected using the “Universal” method optimized for peptide analysis provided by Thermo Scientific. Full MS scans (375–1500 *m/z* range) were acquired in profile mode with the Orbitrap set to a resolution of 120,000 (at 200 *m/z*), cycle time set to 3 seconds and mass range set to “Normal”. The RF lens was set to 30% and the AGC set to “Standard”. Maximum ion accumulation time was set to “Auto”. Monoisotopic peak determination (MIPS) was set to “peptide” and included charge states 2-7. Dynamic exclusion was set to 60s with a mass tolerance of 10ppm and the intensity threshold set to 5.0e3. MS/MS spectra were acquired in a centroid mode using quadrupole isolation window set to 1.6 (*m/z*). Collision-induced fragmentation (CID) energy was set to 35% with an activation time of 10 milliseconds. Peptides were eluted during a 240-minute gradient at a flow rate of 0.250 uL/min containing 2-80% acetonitrile/water as follows: 0-3 minutes at 2%, 3-75 minutes 2-15%, 75-180 minutes at 15-30%, 180-220 minutes at 30-35%, 220-225 minutes at 35-80% 225-230 at 80% and 230-240 at 80-5%.

### C. Label-free quantification (LFQ)

We analysed raw files searched against the Uniprot (www.uniprot.org) *D. melanogaster* database (Dmel_UP000000803.fasta) using Proteome Discover 2.4 (Thermo Scientific). Raw files were searched using SequestHT that included Trypsin as enzyme, maximum missed cleavage site 3, min/max peptide length 6/144, and precursor ion (MS1) mass tolerance set to 20 ppm and fragment mass tolerance set to 0.5 Da and a minimum of 1 peptide identified. Carbamidomethyl (C) was specified as fixed modification, and dynamic modifications set to Aceytl and Met-loss at the N-terminus, and oxidation of Met. A concatenated target/decoy strategy and a false-discovery rate (FDR) set to 1.0% was calculated using Percolator (19). The data was imported into Proteome Discoverer 2.4, and accurate mass and retention time of detected ions (features) using Minora Feature Detector algorithm. The identified Minora features were then used to determine area-under-the-curve (AUC) of the selected ion chromatograms of the aligned features across all runs and relative abundances calculated.

### D. Gene ontology enrichment

We performed gene ontology (GO) enrichment network analyses using the website version of DAVID (v6.8) (20) and Cytoscape (v3.9.0) (21). We used the ClueGO plugin v2.5.8 (22) for Cytoscape to generate enriched GO categories using a right-sided hypergeometric test and *p*-values, adjusted using Benjamini-Hochberg for multiple testing correction, reported. We performed network comparisons between the DmSP2 and DmSP3 using ClueGO. Gene lists were uploaded to DAVID (https://david.ncifcrf.gov/tools.jsp) and functional outputs for all three GO categories (BP, CC, MF) and associated statistical values were saved in Excel spreadsheets. Enriched GO categories with FDR values below 1% are reported. Specific parameters details are found in the figure legends.

### E. Evolutionary rates

We calculated the rate of non-synonymous (dN) to synonymous (dS) nucleotide substitutions (dN/dS) for *D. melanogaster* genes using an existing pipeline (23). We downloaded amino acid sequences and coding sequences (CDS) for D. *melanogaster* (BDGP6.32), and CDS files for *D. sechellia* (dsec_r1.3), *D. simulans* (ASM75419v3), and *D. yakuba* (dyak_caf1) from Ensembl (24). For each species, we identified the longest isoform of each gene and identified orthologs using reciprocal BLASTn (25), with a minimum 30% identity and 1×10-10 E-value cut-off. We identified reciprocal 1:1 orthologs between all four species by the highest BLAST score and identified open reading frames using BLASTx. We then aligned orthologs using PRANK (26) and masked poorly aligned reads with SWAMP (27) using a minimum sequence length = 150, non-synonymous substitution threshold = 7, and window size = 15. We retained 11715 orthologs for analysis after filtering poorly aligned orthologs and those with sequence length < 30bp. We calculated one-ratio estimates (model 0) with an unrooted phylogeny: ((*D. simulans*, *D. sechellia*), *D. melanogaster*, *D. yakuba*), using the CODEML package in PAML (28), and filtered orthologs with a branch specific dS ? 2 or where S*dS ≤ 1 to avoid mutational saturation. In total we retained dN/dS estimates for 11417 genes after filtering, including 2571 (80.9%) proteins in the DmSP3. We tested for differences in evolutionary rates between independent sets of genes using Mann-Whitney U tests.

### F. Experimental design and statistical rationale

We designed experiments to (i) maximize proteome coverage, (ii) measure using label-free quantitation the relative abundance of individual proteins in the proteome and (iii) examine sample purity by measuring the magnitude of adventitious protein binding and contamination in our samples. We performed three independent experiments using three treatments of purified sperm samples as described in Methods. In experiment one, we collected 3 biological replicates of sperm in PBS only. In experiment two sperm were collected in either PBS and Halt protease inhibitor (“Halt” treatment), PBS only (“NoHalt” treatment) or PBS containing 0.1% Triton X100 without protease inhibitor (“PBST” treatment). In experiment three we collected 4 biological replicates of sperm prepared using either PBS (“PBS” treatment) or 2.5M NaCl (“Salt” treatment).

We applied strict thresholds for peptide and protein identification by setting a false-discovery rate (FDR) threshold at 1.0%, calculated using a reverse-concatenated target/decoy strategy in Percolator. To test for differences in abundance between treatments, LFQ ion intensities, calculated using the Minora feature detector in Proteome Discoverer to determine area-under-the-curve (AUC) and summed technical replicates prior to analysis precursor intensities, were fit to protein-wise negative binomial generalized linear models. For experiment two, investigating the effect of detergent treatment, during preliminary analysis we performed pairwise analysis between all three treatments which revealed 16 proteins that showed differential abundance between controls (Halt vs. NoHalt) (Fig. S1). We subsequently performed differential abundance analysis comparing the PBST treatment to the average of both controls (Halt and NoHalt), excluding these 16 proteins (see supplementary analysis). To rank order protein abundances, we calculated a grand mean for each protein, excluding the PBST treatment samples which, as expected, showed substantial differences compared to other samples (see Results).

### G. Statistical analysis

We performed all statistical analysis in R v4.03 (29). All code and analyses are available via GitHub.

To test for non-random distribution of sperm proteins across the polytene chromosomes we downloaded the chromosomal location for all genes in the genome from FlyBase.org (30) and calculated the total numbers of genes on each chromosome. We then summed the observed number of genes found in the sperm proteome on each chromosome, and calculated the expected number based on the total number of sperm proteins identified. We calculated *χ*^2^ statistics for each chromosome and the associated *p*-values with one degree of freedom and used the Benjamini-Hochberg procedure to correct for multiple testing. We excluded analysis of the Y chromosome due to the small number of protein coding genes. To test for non-random distribution of sperm genes across ages classes, we downloaded gene age information from http://gentree.ioz.ac.cn/download.php (31) and grouped as; ancestral (class 0; common to the *Drosophila* genus; n = 12013), subgenus Sophophora (classes 1 + 2; n = 416), melanogaster group (class 3; n = 200), melanogaster subgroup (class 4; n = 334), or recent (classes 5 + 6; n = 120). We tested if sperm genes were randomly distributed across age classes compared to the rest of the genome as above, calculating the observed number of genes in each age class across the genome and among sperm proteins and calculating *χ*^2^ statistics comparing the observed vs. expected number of genes in each age class, using the Benjamini-Hochberg procedure to correct for multiple testing.

To test for differences in abundance between ribosomal proteins compared to the DmSP3 average, independent groups of X-linked-Y-linked- or autosomal-proteins, or between ‘high confidence’ seminal fluid proteins (Sfps), ‘low confidence/transferred’ Sfps, or remaining sperm proteins, we calculated the grand mean abundance across all three experiments excluding the PBST treatment. We filtered proteins identified by two or more unique peptides and found in at least 3 biological replicates in at least 1 treatment group (where applicable). We performed Kruskal-Wallace ranksum tests followed by pairwise Wilcoxon rank-sum tests corrected for multiple testing using the Benjamini-Hochberg procedure. For experiments two and three we performed differential abundance analyses using edgeR (32). For experiment two we filtered proteins with values in 7 out of 9 biological replicates. For experiment three we filtered to include proteins identified in at least 5 replicates (i.e., in at least 3 out of 4 biological replicates of one treatment).

## Results

### H. Overview of the DmSP3

In the current study we identified 2563 proteins across our three experiments (Fig. S2), of which 1965 (76.7%) proteins were identified by two or more unique peptides in a single experiment (n = 1412) or in two or more replicates across any experiment (n = 1867). Relative protein abundances of 2125 proteins (81.2%) were measured by LFQ. As expected from our previous study (33), *α*- and *β*-tubulins, and Sperm-Leucylaminopeptidases (SLaps) were among the most abundant (Table 2). Also present were proteins of unexpected sperm prevalence including ocnus and janus B, a pair of duplicated gene products encoding a testis-specific phosphohistidine phosphatase (34), numerous seminal fluid proteins (Sfps) and over 80 ribosomal proteins (RPs). Overall, we found highly consistent estimates of protein abundances between experiments. Protein abundances were strongly correlated between experiments (Pearson’s correlation = 0.86 - 0.89, all *p* < 0.001; Fig. S3) and median coefficients of variation for each experiment ranged from 0.018-0.054. We performed analyses using the entire DmSP3 (n = 3176; Table S1), combining the 2563 proteins identified in the current study with the 1108 proteins identified in the DmSP2 (7, 14) Fig. 1a).

**Fig. 1.**
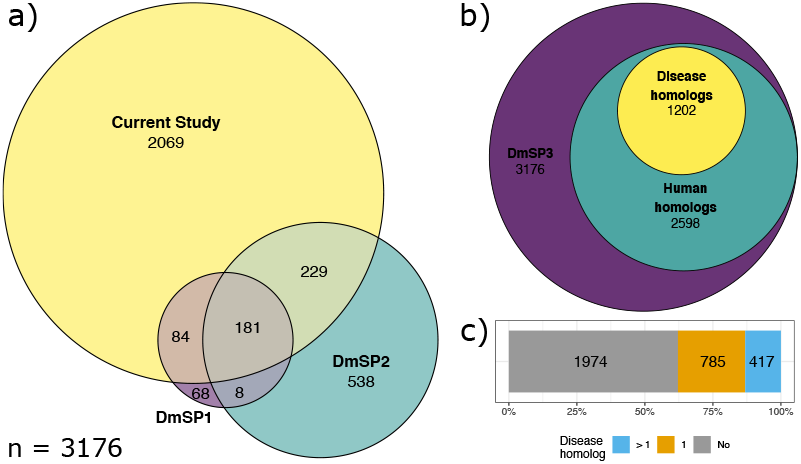
Proteins identified in the *D. melanogaster* sperm proteome (DmSP). a) Overlap between DmSP1, DmSP2 and the current study, together making up the DmSP3 (n = 3176). b) Number of *D. melanogaster* genes found in the DmSP3 with human homologs and disease associated phenotypes from the Online Mendilian Inheritance in Man database (OMIM.org). c) Number of *D. melanogaster* sperm protein genes with none (grey), one (orange), or more than one (blue) associated disease phenotype.

**Table 2.** Most abundant proteins in the DmSP3. Top 20 most abundant proteins in the DmSP3 by LFQ (rank ordered).

| FBgn | Name | Chrom. |
| --- | --- | --- |
| FBgn0003884 | α-Tubulin at 84B | 3R |
| FBgn0003889 | β-Tubulin at 85D | 3R |
| FBgn0259795 | loopin-1 | 2R |
| FBgn0003885 | α-Tubulin at 84D | 3R |
| FBgn0033868 | Sperm-Leucylaminopeptidase 7 | 2R |
| FBgn0035915 | Sperm-Leucylaminopeptidase 1 | 3L |
| FBgn0052064 | Sperm-Leucylaminopeptidase 4 | 3L |
| FBgn0045770 | Sperm-Leucylaminopeptidase 3 | 3L |
| FBgn0039071 | big bubble 8 | 3R |
| FBgn0034132 | Sperm-Leucylaminopeptidase 8 | 2R |
| FBgn0041102 | ocnus | 3R |
| FBgn0031545 | CG3213 | 2L |
| FBgn0037862 | Mitochondrial aconitase 2 | 3R |
| FBgn0038373 | CG4546 | 3R |
| FBgn0002865 | Male-specific RNA 98Ca | 3R |
| FBgn0035240 | CG33791 | 3L |
| FBgn0025111 | Adenine nucleotide translocase 2 | X |
| FBgn0069354 | Porin2 | 2L |
| FBgn0052351 | Sperm-Leucylaminopeptidase 2 | 3L |
| FBgn0012036 | Aldehyde dehydrogenase | 2L |

### I. Gene Ontology and network analyses

The DmSP3 is considerably larger than the DmSP2 (Fig. 1a) and GO analysis identified 24 significantly enriched BP categories (Fig. 2a; Table S2). As expected, major categories included processes involved in energy transduction (e.g., oxidation-reduction, glycolysis, TCA cycle) and reproduction. Other sperm-specific functions included terms related to microtubule and cilium movement. Surprisingly, the GO term “translation” was a prominent member in this analysis containing 78 cytosolic and mitochondrial RPs. To further explore the GO category representation in the DmSP2 and DmSP3, we generated a heat map between the two proteomes in Cytoscape using ClueGO (Fig. 2b). Similar to our previous analysis of the DmSP1 and DmSP2 (14), most of the categories were equal or nearly equal in their shared properties with the one obvious exception being the aforementioned translation BP category as discussed further below.

**Fig. 2.**
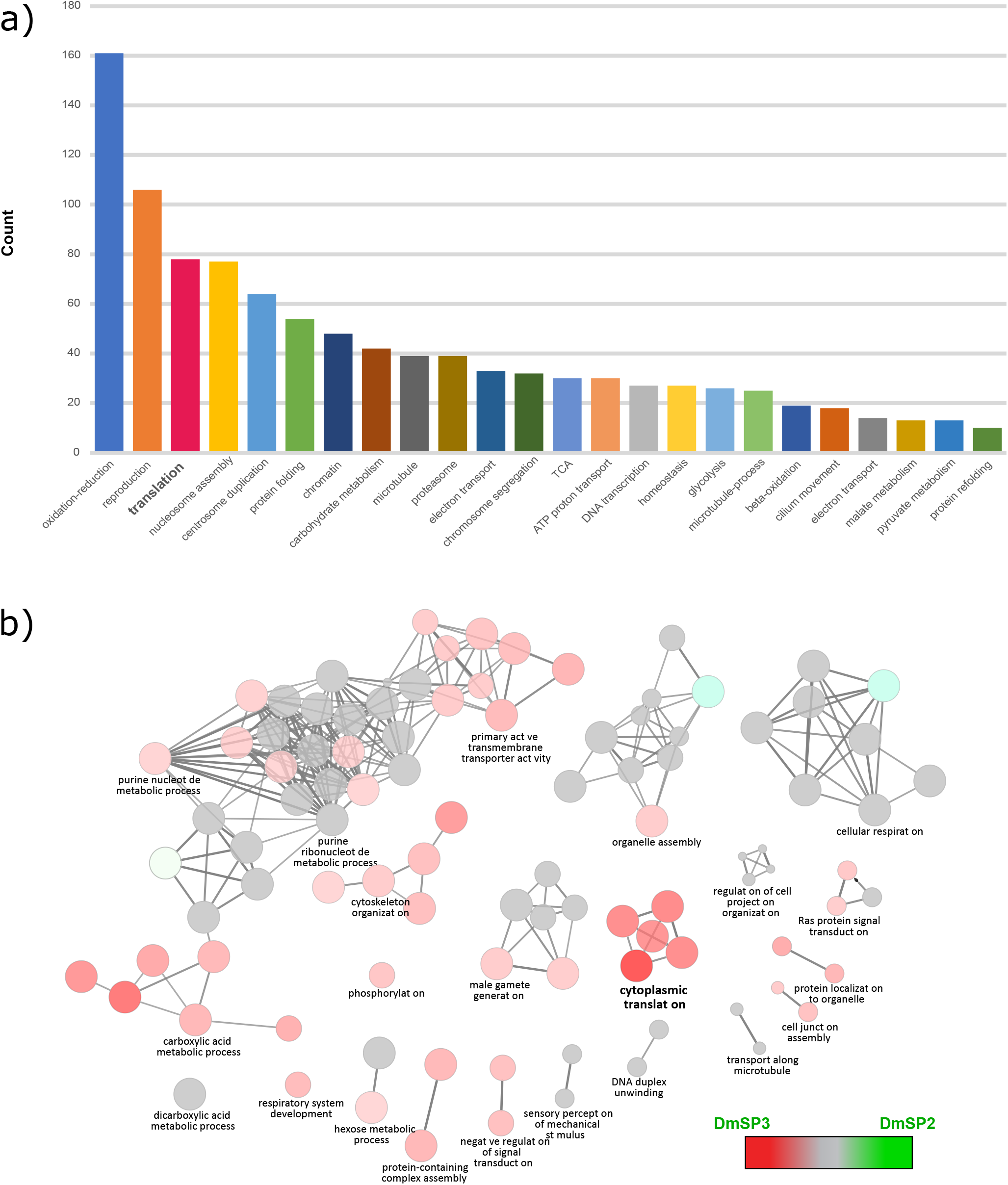
GO functional network enrichment analysis and comparison of the DmSP2 and DmSP3. (a) Bar graph of the 24 GO Biological Process categories identified in the DmSP3 by DAVID (20). Only functional enrichment groups with Benjamini-Hochberg corrected *p*-values < 0.01 and passing a 1% FDR threshold are shown. Note: some GO terms have been combined for clarity; see Table S2 for complete list of GO terms. (b) GO Biological Process network comparison between the DmSP3 (3176 proteins) and DmSP2 (1108 proteins) using the ClueGO plugin for Cytoscape. Color-coded nodes within the network depict the degree of relative compositional enrichment of each dataset. The network is comprised of 22 groups (each comprised of at least 30 genes associated with a common GO functional term) containing a total of 1431 proteins. Node compositional enrichment for proteins identified in the current study (highlighted in red) when node composition bias exceeds 60% while grey nodes indicate equal representation. **Bold** letters indicate one highly enriched category of proteins involved in cytoplasmic translation.

### J. Human disease homologs

Genes in the DmSP3 are highly conserved, with 81.8% (2598/3176) of genes having human homologs, compared to 48% of all *Drosophila* genes (35). Fully 37.8% (1202/3176) of DmSP3 genes have a homolog in humans associated with a known disease or syndrome in a search against the Online Mendelian Inheritance in Man database (OMIM.org; Fig. 1b). Over one third (34.7%; 417/1202) of disease associated DmSP3 genes have more than one human disease homolog (Fig. 1c). Among the most prevalent disease phenotypes found were susceptibility to autism, primary ciliary dyskinesia, spermatogenic failure, and myofibrillar- and congenital-myopathy (Table 3).

**Table 3.** Human disease homologs in the DmSP3. Most common human disease phenotypes from the Online Mendelian Inheritance in Man database (OMIM.org) associated with *D. melanogaster* genes found in the DmSP3. N = number of *D. melanogaster* genes associated with each phenotype. Similar disease phenotypes (marked with an asterisk) have been grouped. Complete list of disease associations can be found in Table S15.

| OMIM phenotype | N |
| --- | --- |
| Autism, Susceptibility To; AUTS20, AUTSX1, AUTSX2 | 27* |
| Ciliary Dyskinesia, Primary; CILD40, CILD3, CILD7 | 25* |
| Spermatogenic Failure; SPGF39, SPGF45, SPGF46 | 24* |
| Myopathy; CFTD, MFM2, Fatal Infantile Hypertonic, Alpha-B Crystallin-Related | 24* |
| Hypertension, Essential | 23 |
| Type 2 Diabetes Mellitus; T2D | 21 |
| Asperger Syndrome, X-Linked, Susceptibility To; ASPGX1, ASPGX2 | 18* |
| Cataract, Multiple Types; CTRCT16, CTRCT9 | 16* |
| Ichthyosis, Congenital, Autosomal Recessive; ARCI4A, ARCI4B | 16* |
| 46, XY Sex Reversal 8; SRXY8 | 12 |
| Colorectal Cancer; CRC | 11 |
| Encephalopathy, Familial, With Neuroserpin Inclusion Bodies; FENIB | 11 |
| Ghosal Hematodiaphyseal Dysplasia; GHDD | 10 |
| Plasminogen Activator Inhibitor-1 Deficiency | 10 |
| Vitamin D-Dependent Rickets, Type 3; VDDR3 | 10 |
| Deafness, Autosomal Recessive 91; DFNB91 | 9 |
| Leukemia, Acute Myeloid; AML | 9 |
| Maturity-Onset Diabetes of The Young, Type 8, With Exocrine Dysfunction; MODY8 | 9 |
| Pseudoxanthoma Elasticum; PXE | 9 |
| Cardiomyopathy, Dilated, 1II; CMD1II | 8 |
| Charcot-Marie-Tooth Disease, Axonal, Type 2F; CMT2F | 8 |
| Neuronopathy, Distal Hereditary Motor, Type IIB; HMN2B | 8 |
| Surfactant Metabolism Dysfunction, Pulmonary, 3; SMDP3 | 8 |

### K. Ribosomal proteins in the DmSP3

Almost one-half of all *D. melanogaster* RPs listed in FlyBase.org (83/169; 49.1%, including paralogs) were identified in the DmSP3 (Table S3). We identified the majority of cytoplasmic RPs (76/93; 81.7%) but only 9.2% (7/76) of mitochondrial RPs. There was no significant difference in RP abundance compared to the DmSP3 average (Kruskal-Wallis rank sum test, *χ*^2^ = 0.063, df = 1, *p* = 0.803; Fig. 3a), suggesting that RPs identified in the DmSP3 are integral to the sperm proteome and not artefactual.

**Fig. 3.**
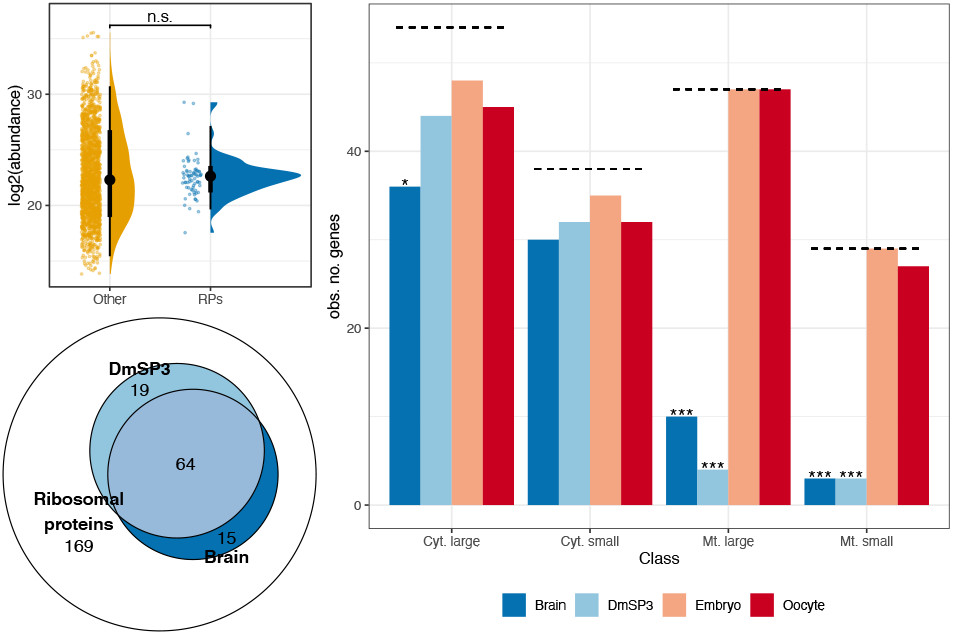
Ribosomal proteins in the DmSP3. a) Abundance of ribosomal proteins (RPs) identified in the DmSP3 compared to the remaining sperm proteome (‘other’). Colored points represent the abundance of individual proteins. Black points show the mean and thick and thin bars represent the 33%, and 66% confidence intervals, respectively. We compared abundances using a Kruskal-Wallace rank sum test. b) Representation of large and small cytoplasmic and mitochondrial RPs in the brain, DmSP3, embryo, or oocyte. The dashed line represents the total number of RPs in each class and asterisks represent results from comparing the observed to expected number of proteins identified using the *χ*^2^ distribution after multiple testing correction. c) Overlap between the total number of RPs identified in the DmSP3 and brain tissue. n.s.; non-significant; * *p* < 0.05; ** *p* < 0.01;*** *p* < 0.001.

The canonical ribosome contains 80 RPs including 13 paralog pairs in *Drosophila* (FlyBase.org). Although the significance of paralog heterogeneity for ribosome function is currently unknown, paralog switching of RPs has been observed in gonads and other tissues (36). We therefore compared RP paralogs in the DmSP3 to those previously described in four tissue types including the testis (36). Significant differences were found between all four tissues as only three RP paralogs were observed in the DmSP3 (RpL22-like, RpS14b and RpS28b) whereas both RPs and paralog RPs were found in all tissues with the exception of two (Rp10Aa and RpS14a; Table S4). Notably, RpL22-like is more abundant in the testis (36), whereas RpL22 was more abundant in the DmSP3. For the remaining paralogs we identified only one member of each pair: the most abundant paralog found in the testis for seven, and the less abundant paralog for two (Table S4). For RpL10Ab and RpS14b only one paralog was identified in both the current study and by Hopes et al. (36), and we did not identify either paralog of RpS10 (RpS10a or RpS10b). Together, these results suggest a complex landscape of paralog switching in the gonad during spermatogenesis and highlight distinct differences between sperm-RP and testis-RP populations.

We next compared the representation of RPs found in the DmSP3 to three other recent proteomic studies in *D. melanogaster* which used Lumos Fusion Orbitrap mass-spectrometers; embryo (37), unfertilised oocyte (38), and brain (39). All four tissue/cell types identified most cytoplasmic RPs, with a slight underrepresentation of large cytoplasmic subunits in the brain (Fig. 3b). The DmSP3 and brain both showed significant underrepresentation of large and small mitochondrial RPs, whereas oocyte and embryo showed almost complete representation of all ribosomal subunits (Fig. 3b). Significantly more RPs identified in the brain or sperm were shared between tissues (64/98; 65.3%) than expected by chance (Fisher’s exact test, *p* < 0.001; Fig. 3c).

### L. Chromosomal distribution of sperm proteins

Sperm proteins were underrepresented on the X-(*χ*^2^ = 12.6, df = 1, *p* = 0.002) and 3L-(*χ*^2^ = 11.8, df = 1, *p* = 0.002) chromosomes (Fig. 4a); a pattern that was previously reported for X-linked genes in the DmSP1 (7) but not replicated in the DmSP2 (14). Protein abundance of X-linked proteins was significantly lower than those on autosomes (Wilcoxon rank-sum test, *p* = 0.041) or the Y chromosome (Wilcoxon rank-sum test, *p* < 0.001; Fig. 4b). We identified 9 of the 16 known proteins encoded on the Y chromosome (Table 4). The average abundance of Y-linked sperm proteins was higher than autosomal sperm proteins (Wilcoxon rank-sum test, *p* < 0.001); 6 within the top 20% most highly abundant proteins, and all within the top 50% (Fig. 4b; Table 4).

**Fig. 4.**
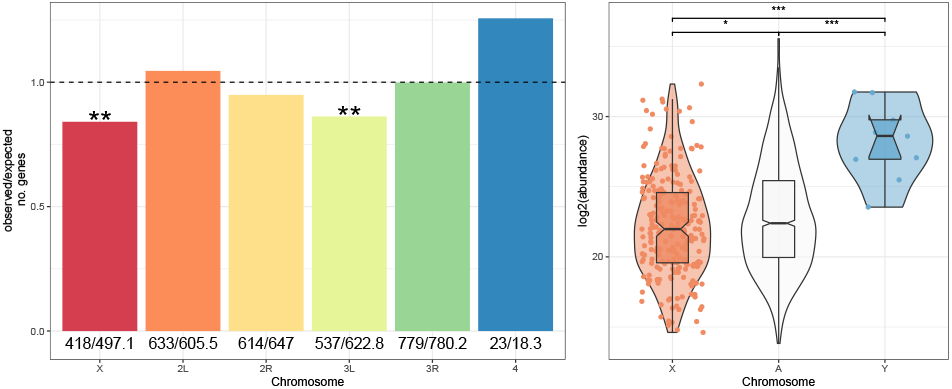
Chromosomal distribution of DmSP3 proteins. a) Chromosomal distribution of sperm proteins. Numbers below bars are the observed and expected number of genes on each chromosome, respectively, and the dashed line indicates the null expectation. Asterisks represent results from comparing the observed to expected number of genes using the *χ*^2^ distribution after multiple testing correction. b) Abundance of sperm proteins found on autosomes (‘A’) and sex chromosomes (‘X’ or ‘Y’). Points, representing individual proteins, are omitted from autosomes for clarity. Asterisks represent results from pairwise Wilcoxen rank-sum test corrected for multiple testing using the Benjamini-Hochberg procedure. n.s., non-significant; * *p* < 0.05; ** *p* < 0.01; *** *p* < 0.001.

**Table 4.** Y-linked sperm proteins in the DmSP3. Genes are rank ordered by mean abundance and link to male fertility from gene knockout/knockdown experiments (40, 41) are shown.

| FBgn | Name | Ranked abundance (%) | Sterile |
| --- | --- | --- | --- |
| FBgn0267433 | male fertility factor kl5 | 98.8 | Yes |
| FBgn0267432 | male fertility factor kl3 | 98.8 | Yes |
| FBgn0058064 | Aldehyde reductase Y | 95.5 | No |
| FBgn0001313 | male fertility factor kl2 | 93.6 | Yes |
| FBgn0046323 | Occludin-Related Y | 92.9 | No |
| FBgn0267449 | WD40 Y | 86.4 | Yes |
| FBgn0267592 | Coiled-Coils Y | 86 | Not studied |
| FBgn0046697 | Ppr-Y | 78.3 | No |
| FBgn0046698 | Protein phosphatase 1, Y-linked 2 | 65.2 | No |

### M. Seminal fluid proteins identified in the DmSP3

Sfps have been extensively studied in *Drosophila* with over 600 putative Sfps identified to date including 292 that are considered ‘high confidence’ (42). A surprisingly high number of Sfps were identified in the DmSP3 (122 ‘high confidence’ Sfps; 156 ‘low confidence/transferred’ Sfps; Table S1) (42).

We found no significant difference in abundance between Sfps and the remaining DmSP3 (Kruskal-Wallace rank-sum test, *χ*^2^ = 4.28, df = 1, *p* = 0.118; Fig. 5a) and 44 ‘high confidence’ Sfps were at, or above, the median abundance of the DmSP3 (Table S5).

**Fig. 5.**
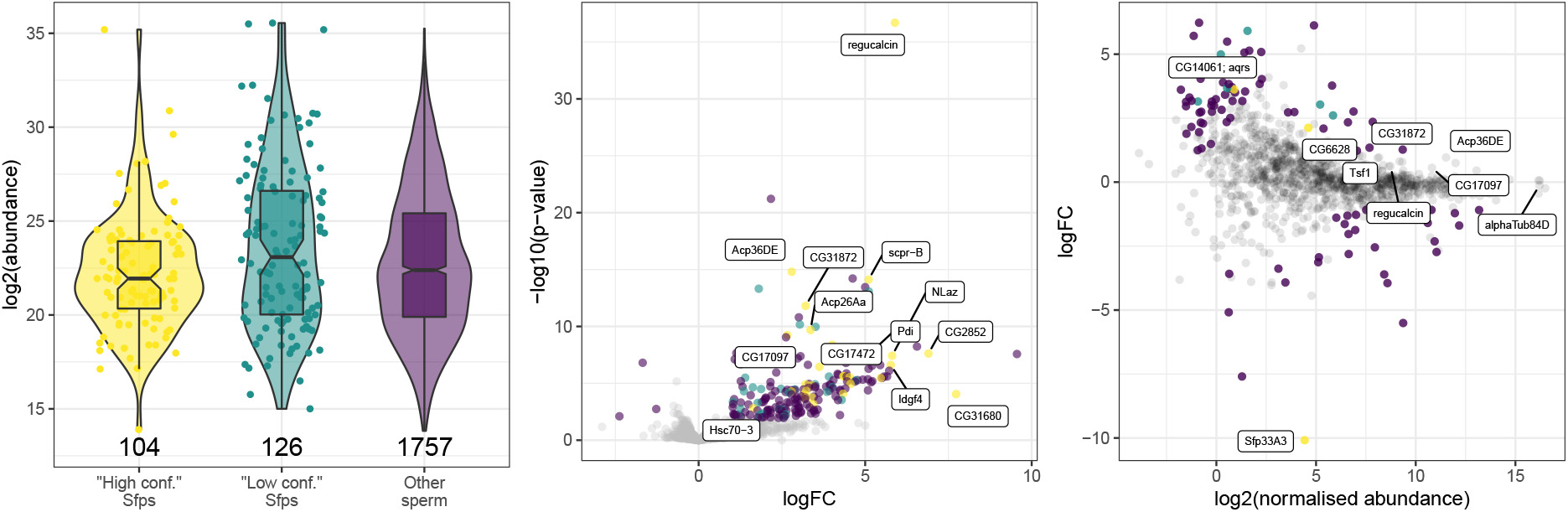
Seminal fluid proteins in the DmSP3. a) log2 abundance of proteins found in the DmSP3 classified as ‘high confidence’ Sfps, ‘low confidence or transferred’ Sfps by Wigby et al. (42), or remaining sperm proteins. Points, representing individual proteins, are omitted from ‘other sperm’ for clarity. b) Volcano plot for difference between PBST treatment vs. the average of both controls (Halt and NoHalt) in experiment two. Positive values indicate higher abundance in controls. c) MA plot for difference between NaCl treatment vs. PBS control. Positive values indicate higher abundance in NaCl treatment. For b) and c) points are coloured as in a) denoting ‘high confidence’ (yellow) and ‘low confidence/transferred’ (turquoise) Sfps or remaining sperm proteins (purple) that showed significant differences in abundance based on a |logFC| > 1 and false discovery rate corrected *p*-value < 0.05. Several Sfps are labelled in b) that showed differential abundance between treatments. Sfps are labelled in c) that were among the top 10% most abundant proteins, and the three Sfps that showed significant differences in abundance between treatments (Sfp33A3, aquarius [CG14061], and CG6628).

We therefore examined the binding characteristics of the Sfps by washing purified sperm with a strong anionic detergent (Triton X-100) known to disrupt plasma membranes. Following detergent treatment 1600 proteins were identified, the majority (1063/1600; 66%) identified by 2 or more unique peptides. We identified 198 proteins that were lower abundance in PBST compared to controls (Table S6) and three proteins more abundant in PBST samples (Fig. 5b).

Of the 60 ‘high confidence’ Sfps identified by two or more unique peptides in experiment two, 17 (28.4%) were filtered out prior to analysis (including 14 which were not detected in PBST samples in any replicate), and 29 (48.3%) were found at significantly lower abundance in PBST samples, together suggesting these proteins are weakly bound or found on the sperm plasma membrane. The remaining 14 (23.3%) Sfps showed no significant difference in abundance, suggesting tight association with sperm (Table 5). Additionally, 13 out of 53 (24.5%) RPs detected in experiment two were significantly lower in abundance after PBST treatment. Proteins lower in abundance after PBST treatment showed GO enrichment of multicellular organism reproduction, mitochondrial transport, transmembrane transport, cytoplasmic translation, and sarcomere organisation (BP). Thus, as expected, PBST treatment stripped lipids and membrane- and membrane-bound proteins (including Sfps) from sperm (Table S7).

**Table 5.** Seminal fluid proteins remaining in the sperm proteome after PBST treatment.

| FBgn | Name | Chrom. |
| --- | --- | --- |
| FBgn0011694 | Ejaculatory bulb protein II | 2R |
| FBgn0261055 | Seminal fluid protein 26Ad | 2L |
| FBgn0004181 | Ejaculatory bulb protein | 2R |
| FBgn0003885 | α-Tubulin at 84D | 3R |
| FBgn0260745 | midline fasciclin | 3R |
| FBgn0036970 | Serpin 77Bc | 3L |
| FBgn0036969 | Serpin 77Bb | 3L |
| FBgn0259975 | Seminal fluid protein 87B | 3R |
| FBgn0034709 | Secreted Wg-interacting molecule | 2R |
| FBgn0264815 | Phosphodiesterase 1c | 2L |
| FBgn0020414 | Imaginal disc growth factor 3 | 2L |
| FBgn0050104 | Ecto-5'-nucleotidase 2 | 2R |
| FBgn0052203 | Serpin 75F | 3L |
| FBgn0003748 | Trehalase | 2R |

In experiment three, we washed sperm samples with high molar salt expected to weaken ionic bonds and eliminate non-specific protein binding to sperm (including Sfps). We identified 1890 proteins, of which 1273 (65%) were identified by two or more unique peptides. After filtering (see Methods) we performed differential abundance analysis for 1202 proteins and identified 92 differentially abundant proteins, including 3 Sfps (Sfp33A3, aquarius [CG14061], and CG6628) (Fig. 5c). The remaining 48 ‘high confidence’ Sfps we identified in this experiment did not show significant differential abundance between treatments, with 6 Sfps in the top 20% most abundance proteins (regucalcin, Acp36DE, CG31872, Transferrin 1, CG17097, and *α*-Tubulin at 84D).

### N. Gene – protein abundance concordance

To explore the relationship between protein abundance and gene expression for the 68 ‘high confidence’ Sfps tightly binding to sperm following detergent or salt treatment (‘sperm associated Sfps’; Table S8), we compared gene expression (FPKM; fragments per kilobase of transcript per million mapped reads) for all proteins identified in the DmSP3 between the accessory glands, carcass, ovary, and testis using data retrieved from FlyAtlas2 (43). The average expression of both sperm associated Sfps and the remaining Sfps identified in the DmSP3 was highest in the accessory glands, while the remaining DmSP3 proteins were most highly expressed in testis (Fig. S4a). However, 7 sperm associated Sfps showed higher expression in the testis than accessory glands (Table S9).

The abundance of proteins in the DmSP3 had the strongest correlation (*β*) and best fit (*R*^2^) in the testis (*β* = 0.460, *R*^2^ = 0.133, *p* < 0.001, n = 1498) (Fig. S4b). Protein abundance of sperm associated Sfps was positively correlated with gene expression in the testis (*β* = 0.399, *R*^2^ = 0.152, *p* = 0.006, n = 49), but not the accessory glands (*p* = 0.246), carcass (*p* = 0.052), or ovary (*p* = 0.271). The abundance of remaining Sfps identified in the DmSP3 was positively correlated with gene expression in the accessory glands (*β* = 0.274, *R*^2^ = 0.197, *p* = 0.004, n = 41) and testis (*β* = 0.281, *R*^2^ = 0.147, *p* = 0.040, n = 29), but not the carcass (*p* = 0.109) or ovary (*p* = 0.677) (Fig. S4c). Therefore, our results suggest sperm associated Sfps show tighter regulation with gene expression in the testis than accessory glands.

### O. Gene age

A variety of mechanisms drive genomic and protein diversity including gene duplication and retroposition (6, 31) resulting in unique, lineage-specific patterns of gene age (44–46). Newly evolved genes frequently acquire testis-biased gene expression (47) and it was therefore of interest to query the gene age landscape of the DmSP3. There were fewer ‘recent’ (*χ*^2^ = 6.58, df = 1, *p* = 0.026), ‘melanogaster subgroup’ (*χ*^2^ = 9.69, df = 1, *p* = 0.009), and ‘Sophophora-group’ (*χ*^2^ = 5.51, df = 1, *p* = 0.032) age genes than expected by chance, indicating genes encoding sperm proteins are underrepresented in more recent evolutionary time (Fig. 6a). We identified 13 genes of recent origin, of which five were located on the X chromosome (Table S10).

**Fig. 6.**
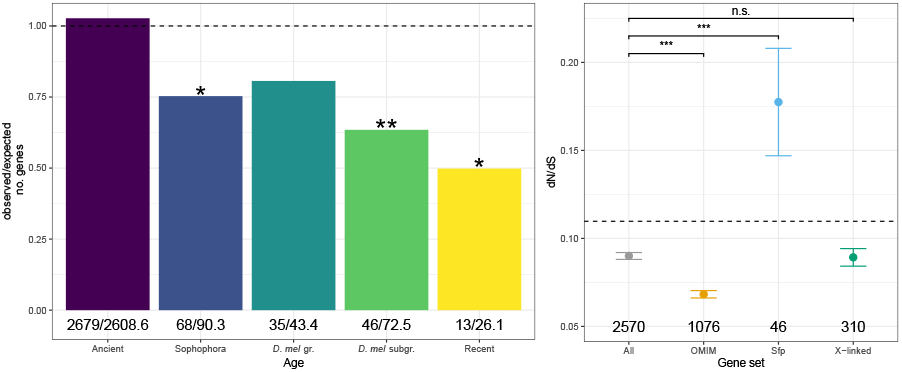
Sperm evolutionary dynamics. a) Gene age distribution of sperm proteins. Numbers below bars are the observed and expected number of genes in each age class, respectively, and the dashed line at 1 indicates the null expectation. Asterisks represent results from comparing the observed to expected number of genes using the *χ*^2^ distribution after multiple testing correction. b) Mean (± standard error) nonsynonymous (dN) to synonymous (dS) nucleotide substitution rate (dN/dS) estimates for sperm proteins. Asterisks represent results from Mann-Whitney U tests comparing each gene set (OMIM, Sfp, X-linked) to the genome average (“All”), excluding proteins in that set. Dashed line represents the genome average (mean dN/dS = 0.110, standard error = 0.001, n = 11417). Numbers below points indicate numbers of genes in each category. Note: groups are not necessarily mutually exclusive, i.e., ‘OMIM’ proteins may also be ‘X-linked’, etc. n.s., non-significant; * *p* < 0.05;** *p* <0.01;*** *p* <0.001.

### P. Sperm evolutionary rates

Genes in the DmSP3 evolve more slowly than the genome average (Mann-Whitney U test, *p* < 0.001). This pattern remains when considering X-linked sperm proteins compared to the genome average (*p* < 0.001), which evolve at a similar rate to the DmSP3 average (*p* = 0.958; Fig. 6b). Sfps in the DmSP3 evolve faster than the DmSP3 average (*p* < 0.001), at a similar rate to other Sfps (*p* = 0.232; Fig. S5), whereas genes with a human disease homolog (OMIM.org) evolve more slowly than the DmSP3 average (*p* < 0.001; Fig. 6b).

The top 10%, fastest evolving genes in the DmSP3 (dN/dS [mean ± s.e.] = 0.313 ± 0.009, n = 257, Table S11) showed GO enrichment for multicellular organism reproduction (BP) and extracellular space (CC) (Table S12). The bottom 10%, slowest evolving genes in the DmSP3 (dN/dS = 0.004 ± 0.0002, n = 258, Table S13) showed GO BP enrichment for cytoplasmic translation, centrosome duplication, regulation of cell shape, ribosomal large subunit assembly, tricarboxylic acid cycle, ATP hydrolysis coupled proton transport, cell adhesion, oocyte microtubule cytoskeleton polarisation, and endocytosis (Table S14).

## Discussion

In summary, our reanalysis of the *D. melanogaster* sperm proteome (DmSP3) more than doubled the number of identified proteins, dramatically increased representation of RPs, and highlighted several human neurological disease homologs. LFQ identified highly abundant tubulins, Sperm-Leucylaminopeptidases (S-Laps), Y-linked sperm proteins and ocnus, a testis-specific protein. LFQ also provided direct evidence for lowered abundances of X-linked sperm proteins. Sperm genes evolve relatively slowly and are underrepresented in recent age classes, consistent with evolutionary constraint acting on the sperm proteome. Finally, we identified a number of Sfps in the DmSP3 which were resistant to detergent or high molar salt treatment, suggesting some are integral to the sperm proteome.

The increased (> 2-fold) depth of proteome coverage is likely due to improved protein extraction, efficiency of trypsination/peptide recovery and direct injection methods employed in this study. Traditionally SDS-PAGE off-line pre-fractionation has been the method of choice for the analysis of complex proteomes. However, these off-line methods come at a cost: sample loss due to the extra steps involved and the well-known issues of peptide recovery from polyacrylamide gels (48, 49). Although work to alleviate this limitation continues to improve this approach, our results suggest that a combination of high SDS concentrations in the initial solubilization and use of immobilized enzymatic digestion using S-Trap technologies greatly enhanced the yield of usable peptides for bottom-up proteomics. The DmSP3 also contained Yolk protein 2 (Yp2), a protein previously found in sperm (50) but undetected in the DmSP1 or DmSP2. As noted by the authors of this study, detection of Yp2 in sperm required large amounts of input protein for detection on immunoblots, suggesting Yp2 was present at very low levels in the testis and sperm (50). Therefore, detection of Yp2 in our study provides additional confidence in the efficacy or our approach.

The sperm proteome is expected to exhibit dynamic gene movement and expression evolution due to its sex-specific expression and essential role for male fertility (6). We found X-linked genes are underrepresented in the sperm proteome, as reported in the DmSP1 (7). Additionally, we show that X-linked sperm proteins were found in significantly lower abundances, consistent with the downstream effects of meiotic sex chromosome inactivation (16–18), and/or resolution of intralocus sexual conflict (51–53). In contrast, more than half of Y-linked proteins (9/16) including known fertility factors were present in the DmSP3 (40, 41). Our LFQ analysis revealed all 9 Y-linked protein abundances were above the DmSP average, with 7/9 in the top 10%. This is the first quantitative assessment of this important class of sperm proteins in sperm and adds direct empirical evidence in support of the long-standing hypothesized structural role in the assembly of the sperm axoneme (54).

We found sperm proteins evolve more slowly than the genome average. Slow rates of adaptive evolution could be due to purifying selection or weak selection acting on sperm genes as they are shielded from selection in females (7, 55, 56). Sperm proteins were also underrepresented in recent evolutionary age classes and over 80% had human homologs, supporting the idea that sperm genes are under evolutionary constraint. A recent study found Sfps are overrepresented in recent age classes (57), indicating different evolutionary forces acting on sperm vs. non-sperm components of the ejaculate. Sfps in the DmSP3 evolve at a similar rate to Sfps found elsewhere in the genome, and more quickly than the DmSP3 average, suggesting similar evolutionary pressures affecting rates of Sfp evolution across tissue types.

The abundance of RPs in the DmSP3 was unexpected given that sperm are stripped of most cellular machinery prior to maturation. However, sperm may undergo post ejaculatory modifications, perform secondary sexual functions, or provision the developing zygote after fertilization, requiring protein synthesis (58–61). Sperm function beyond delivering a haploid compliment of nuclear material for fertilization still remains relatively underexplored (59, 62, 63). The presence of a large repertoire of core RPs delivered to the egg during fertilization raises the intriguing possibility that paternally-derived ribosomes are active during zygote formation and perhaps beyond.

Another intriguing finding that sperm had higher abundance of RpL22 versus the paralog RpL22-like, opposite from levels found in the testis (36) suggests a complex pattern of paralog switching and selectivity during spermatogenesis. While the functional significance of this selectivity is unknown, they are interesting to consider in the context of the known mRNA repertoire in *Drosophila* sperm delivered to the egg at fertilization (61). Fully 33% of the total sperm mRNA repertoire encoded ribosomal proteins (47/142; ref. (61)), a striking coincidence that warrants further study. We also found similarity in the underrepresentation of mitochondrial RPs in both the DmSP3 and brain, providing yet another example of the molecular similarities between these two tissue types (64). Finally, we note that the DmSP3 contains as many as 300 entries with GO annotation terms related to neuronal structure and function, lending additional support to the similarities drawn between the brain and testis.

### Q. Possible testis origin of seminal fluid proteins

Although some Sfps were previously identified, but not quantified, in the DmSP2 (14), the unexpectedly high numbers (and in some cases, relative abundances) of Sfps in the DmSP3 adds to an expanding landscape of seminal fluid protein biology. As Sfps are thought to be primarily secreted from the paired accessory glands and the ejaculatory bulb in *Drosophila* (65), our results raise the possibility that some Sfps are integral to the sperm proteome, are secreted from the testes or seminal vesicles, or bind to sperm prior to mixing in the ejaculatory duct. We identified 122 ‘high confidence’ Sfps (42) in the DmSP3 which is unlikely artefactual given that many Sfps were found in multiple biological replicates and in independent experiments. Denaturing the sperm plasma membrane using detergent stripped most (75%) Sfps from the sperm proteome, suggesting these Sfps are integral to the sperm plasma membrane or bound to sperm advantageously in the seminal vesicles prior to mixing in the ejaculatory duct. High molar salt had little effect on the composition of the sperm proteome, indicating some Sfps are bound strongly to sperm.

We identified 68 ‘sperm associated Sfps’ that were not depleted by detergent or salt treatment. We suggest several of the ‘high-confidence’ Sfps identified in the DmSP3 that are highly expressed in the testes (Table S9) should be classified as sperm proteins. In addition, *α*-Tubulin at 84D (FBgn0003885) is a major constituent of microtubules and involved in sperm axoneme assembly, and therefore likely a sperm protein. Notably, Acp36DE was consistently among the most abundant proteins in our experiments. Acp36DE tightly binds sperm and is essential for efficient sperm storage in the female sperm storage organs (66, 67). The possibility that Sfps bind to sperm in the seminal vesicles prior to mixing in the ejaculatory duct should be investigated further. Moreover, the potential for the testes, seminal vesicles, or perhaps even sperm cells, to secrete proteins, including Sfps, requires further investigation.

Finally, the DmSP3 contains over 1200 human disease homologs. The prominence of several neurological diseases (e.g., Primary Ciliary Dyskinesia, susceptibility to autism, encephalopathy, and neuropathy) may be related to the shared functional designs of sperm and neurons, cells of extraordinary axial ratios transmitting biological information over large distances. It will be of great interest to tease out the significance of this subset of neural-related DmSP3 proteins in the context of sperm function and its related reproductive activities and possible relevance for study of human diseases.

### R. Conclusion

Our reanalysis of the *D. melanogaster* sperm proteome using improved separation and detection methods and an updated genome annotation highlights several key features of sperm function and evolution, including the prominence of proteins integral to sperm development (tubulins and S-Laps), the dynamic nature of sex-linked sperm genes, and constraints on sperm proteome evolution. We also show the prevalence of many RPs, despite the expectation that sperm are transcriptionally silent. The parallels in ribosomal protein composition and occurrence of several human neurological disease homologs also lends further support to the functional similarities between sperm and neurons. Finally, we demonstrate that a significant number of seminal fluid proteins are found in the sperm proteome raising the possibility that Sfps mix with sperm in the seminal vesicles, or Sfps may be secreted from the testes, seminal vesicles, or even sperm cells.

## Abbreviations

AUC: area-under-the-curve
BP: biological processes
CC: cellular components
CDS: coding sequences
CID: collision-induced fragmentation
DAVID: database for visualisation and integrated discovery
DmSP: *Drosophila melanogaster* sperm proteome
FBgn: FlyBase gene number
FDR: false discovery rate
FPKM: fragments per kilobase of transcript per million mapped reads
GO: Gene ontology
LFQ: Label-free quantitation
MF: molecular functions
MIPS: monoisotopic peak determination
OMIM: Online Mendelian Inheritance in Man
PAML: phylogenetic analysis by maximum likelihood
RPs: ribosomal proteins
S-Laps: sperm leucyl-aminopeptidases
Sfp: Seminal fluid protein
TEAB: triethylammonium bicarbonate

## S. Data availability

Proteomic data have been deposited to the [ProteomeXchange Consortium](http://proteomecentral.proteomexchange.org/) via the PRIDE partner repository (68) with the identified PDXXXXXXXX. All code and analyses are available on [GitHub](https://martingarlovsky.github.io/DmSP3/).

## T. Author contributions

TLK and MDG conceived the study. TLK and JS performed dissections and LC-MS experiments. MDG and TLK performed analyses and wrote the manuscript. All authors agreed on the final version of the manuscript.

## ACKNOWLEDGEMENTS

We would like to thank Caitlin McDonough-Goldstein and Maria Vibranovski for helpful discussion, Alison Wright, Daniela Palmer, and Leeban Yusef for advice analysing evolutionary rates, and Eric Sedore and Larne Pekowsky from the Syracuse University HTC Campus Grid and NSF award ACI-1341006 for providing com-puting services. We are also grateful to the authors whose data was used in this study for making data publicly available and the curators of FlyBase.org for continued maintenance of this essential resource. This work was funded in part by the Biodesign Institute and ASU Knowledge Enterprise Core Research Facilities.

## Supplementary Note 1: Supplementary information

**Fig. S1.**
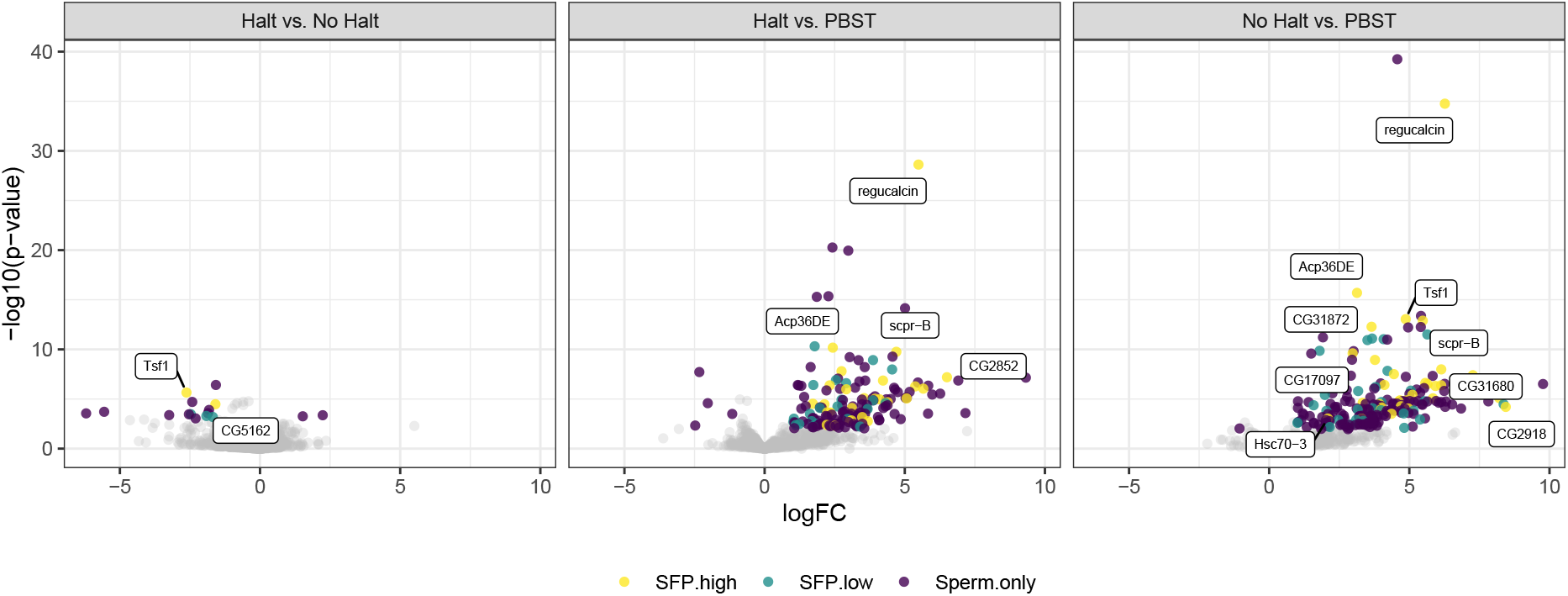
Volcano plots from pairwise analyses between treatments in experiment two, denoting ‘high confidence’ (yellow) and ‘low confidence/transferred’ (turquoise) Sfps or remaining sperm proteins (purple) that showed significant differences in abundance based on a |logFC| > 1 and false discovery rate corrected *p*-value < 0.05. Several Sfps are labelled that showed differential abundance between treatments.

**Fig. S2.**
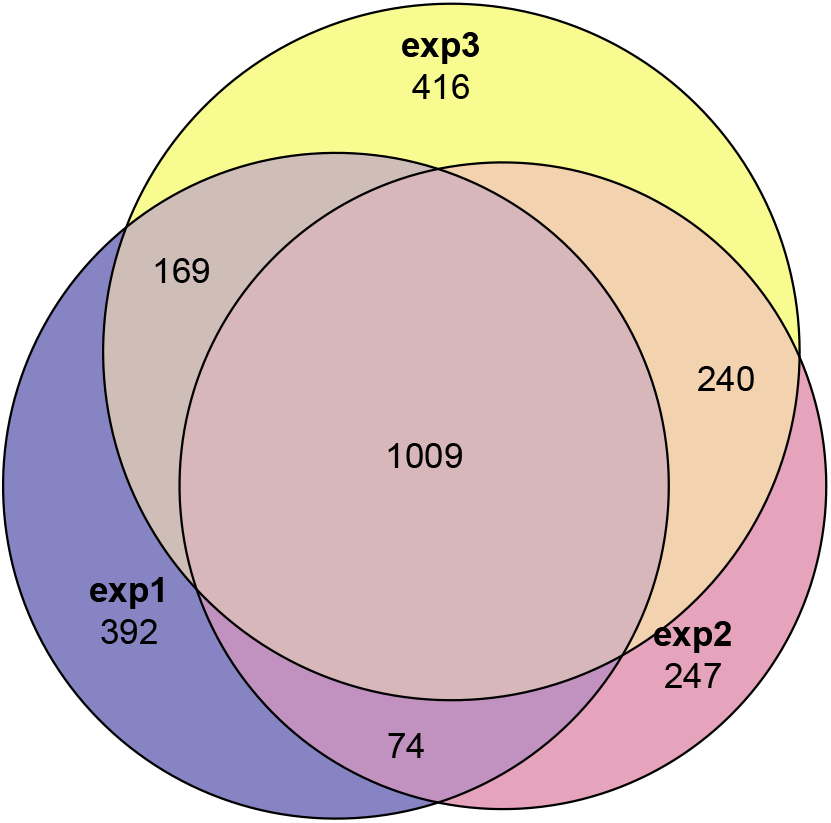
Overlap between proteins identified in each experiment in the current study.

**Fig. S3.**
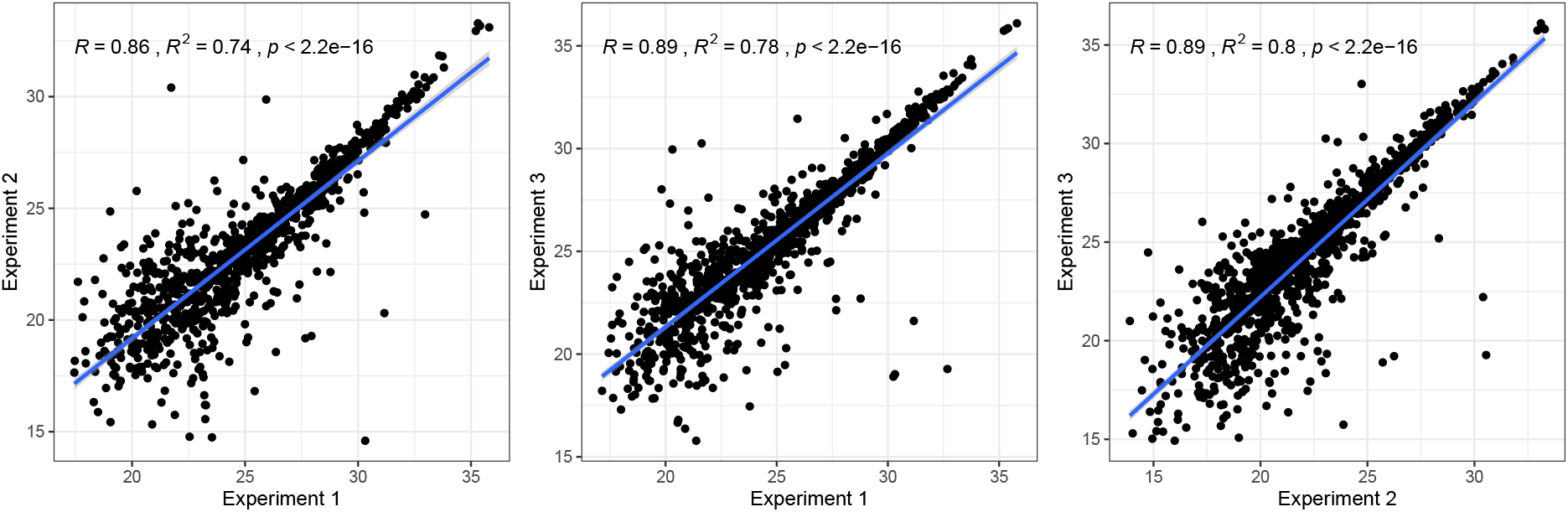
Correlations in average protein abundance between each experiment in the current study. Mean protein abundance was calculated across all replicates for each experiment, except experiment two which excluded the PBST treatment. Shown are Pearson’s correlations and line of best fit.

**Fig. S4.**
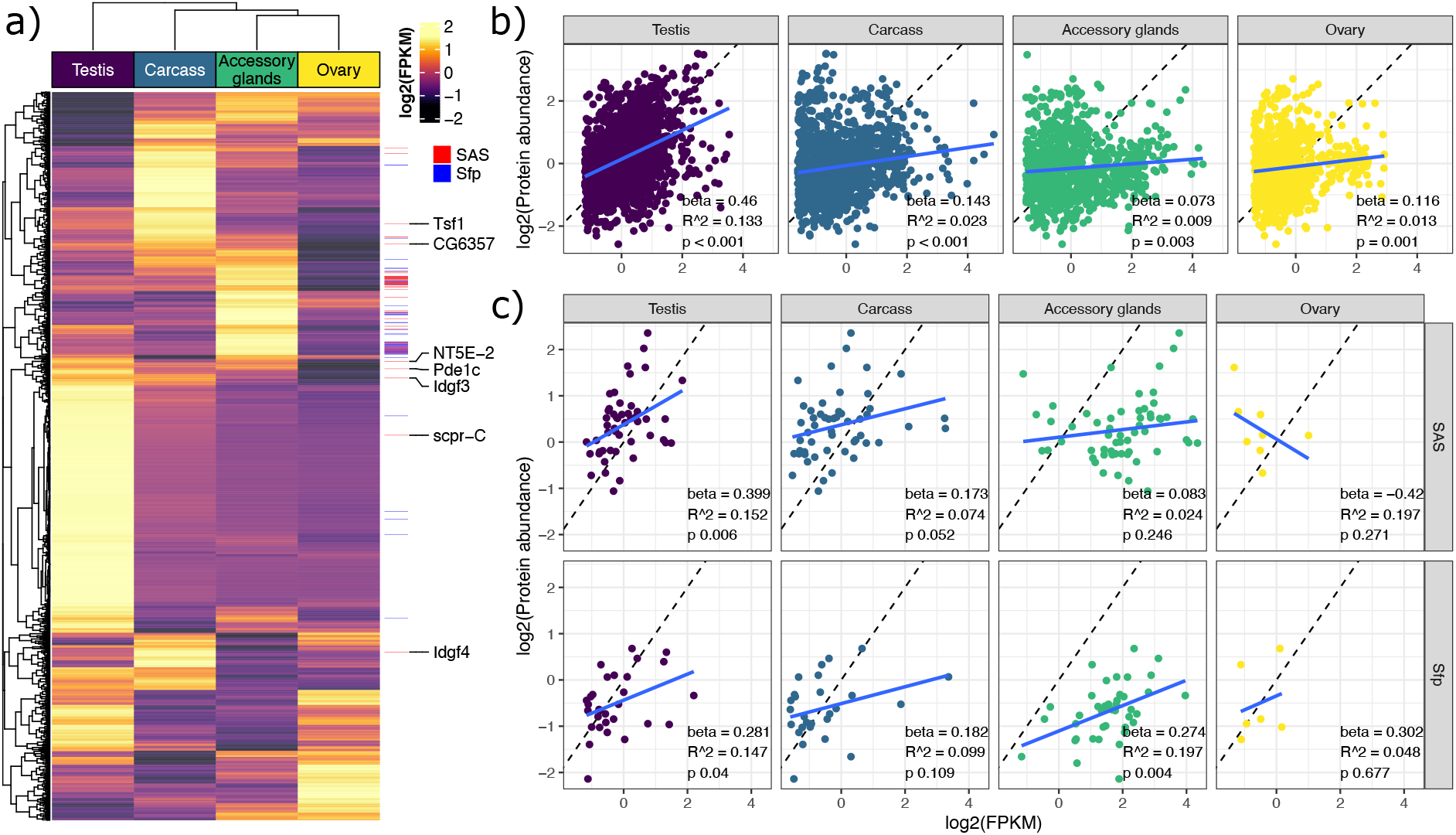
Gene – protein abundance concordance in the DmSP3. a) Heatmap of mRNA expression of DmSP3 genes (n = 2673) in the accessory glands, carcass, ovary, and testis. Data retrieved from from FlyAtlas2 (43) are log2(FPKM) scaled per gene. The 7 ‘high confidence’ Sfps with higher expression in the testis than accessory glands are highlighted on the right. Labels on the right also show ‘sperm associated Sfps’ (red) and other ‘high confidence’ Sfps (blue) identified in the DmSP3. b) Linear regressions of gene expression on protein abundance in the testis (n = 1498), carcass (n = 1165), accessory glands (n = 1001), and ovary (n = 825). c) Linear regressions of gene expression on protein abundance for ‘sperm associated Sfps’ (SAS) and remaining Sfps identified in the DmSP in each tissue. b) and c) are linear regressions using z-score log2-transformed values after filtering genes with log2-FPKM < 2. Blue lines are model fits from a linear regression, dashed lines indicate a perfect correlation between gene expression and protein abundance.

**Fig. S5.**
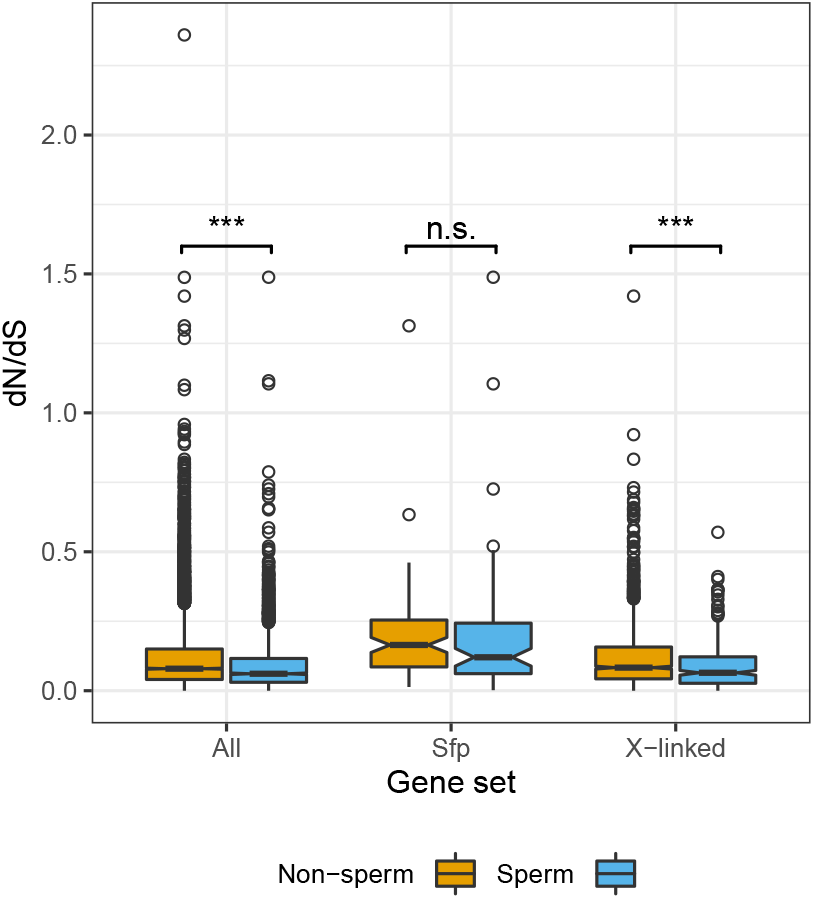
Nonsynonymous (dN) to synonymous (dS) nucleotide substitution rate (dN/dS) estimates for proteins in the DmSP3 or elsewhere. Asterisks represent results from Mann-Whitney U tests; n.s., non-significant; ***, *p* < 0.001.

## Notes

https://martingarlovsky.github.io/DmSP3/

### Competing Interest Statement

The authors have declared no competing interest.

